# *Enterobacter* sp. SA187 mediates plant thermotolerance by chromatin modification of heat stress genes

**DOI:** 10.1101/2020.01.16.908756

**Authors:** Kirti Shekhawat, Arsheed Sheikh, Kiruthiga Mariappan, Rewaa Jalal, Heribert Hirt

## Abstract

Global warming has become a critical challenge to food safety, causing severe yield losses of major crops worldwide. Heat acclimation empowers plants to survive under extreme temperature conditions but the potential of beneficial microbes to make plants thermotolerant has not been considered so far. Here, we report that the endophytic bacterium *Enterobacter* sp. SA187 induces heat tolerance in *Arabidopsis thaliana* by reprogramming the plant transcriptome to a similar extent as acclimation. Acclimation induces priming of heat stress memory genes such as *APX2* and *HSP18.2* via the transcription factors *HSFA1A, B, D, and E* and the downstream master regulator *HSFA2. hsfa1a,b,d,e* and *hsfa2* mutants compromised both acclimation and bacterial priming through the same pathway of *HSF* transcription factors. However, while acclimation transiently modifies H3K4me3 levels at heat stress memory gene loci, SA187 induces the constitutive priming of these loci. In summary, we demonstrate the molecular mechanism by which SA187 imparts thermotolerance in *A. thaliana*, suggesting that beneficial microbes might be a promising way to enhance crop production under global warming conditions.

## Introduction

Global warming increases average temperature and causes severe heat waves, challenging plant growth and agriculture yield worldwide (Mittler et al, 2010; Ling et al, 2018). Elevated temperatures cause severe cellular injury to plants resulting in a collapse of cellular organization and inhibition of plant growth (El-Daim et al, 2014). In order to cope with heat stress, plants have developed several strategies such as basal heat tolerance and acquired heat tolerance. In basal heat tolerance, plants have a natural capacity to deal with heat stress, whereas in acquired thermotolerance, plants acquire tolerance to lethal levels via a short pre-exposure to mild heat stress, a phenomenon that is known as priming (Yeh et al, 2012). In priming, plants establish a molecular stress memory of previous exposure to mild heat stress (Sani et al, 2013). This molecular memory is responsible for the higher expression of the heat stress transcription factors (*HSF*s) regulating expression of heat shock proteins (*HSP*s) and antioxidant genes. *HSP*s act as molecular chaperones that protect the conformation and function of proteins under heat stress. Therefore, stress memory allows primed plants to respond robustly and quickly to subsequent exposure to heat stress, which helps their recovery (Prime-A-Plant Group, 2006; Hilker et al, 2016; Lin et al, 2016). HEAT SHOCK FACTOR A2 (HSFA2) is required for an active heat stress memory in plants (Charng et al, 2007). Among 21 other Heat Shock Factors, *HSFA2* is known to be the only *HSF* gene that has a specific function in heat stress memory in *Arabidopsis thaliana* (Scharf et al, 2012; Ohama et al, 2017). The expression of *HSFA2* is further regulated by four isomers of *HSFA1A, B, D* and *E*, which are considered to be master regulators of heat stress because they activate the expression of *HSFA2*. In turn, *HSFA2*, amplifies the transcriptional induction of a subset of heat stress response genes (Schramm et al, 2008; Liu et al, 2011; Yoshida et al, 2011; Yeh et al, 2012, Mishra et al, 2002; Liu et al, 2011; Yoshida et al, 2011; Liu and Charng, 2013). The subset of genes, to which *HSFA2* binds to, is known as memory genes or heat stress responsive genes because of their persistent transcriptional induction after heat stress (Stief et al, 2014). *HSFA2* binds transiently in a hit and run mode at the promoter region of memory genes. This binding facilitates the chromatin modifications that are responsible for longer induction of heat stress memory genes (Lamke et al, 2016). The di- and trimethylation of lysine 4 on histone H3 (H3K4me2, H3K4me3) showed a lasting enrichment at the *APX2* and *HSP18.2* memory genes and persist up to 5 days after the priming heat stress. Therefore, heat acclimation induces a transcriptional memory that also involves chromatin modifications and leads to a hyper transcriptional response upon re-occurring heat stress (Bruce et al, 2007; Vriet et al, 2015; Baurle, 2017; Lamke and Baurle 2017).

Heat priming could be applied in agriculture to make crops more stress-resistant and productive (Liu and Charng 2012). However, acclimatizing plants under field conditions is not always feasible. The use of endophytes and other rhizobacteria (known as plant growth promoting bacteria, PGPB) as a reliable and feasible method to promote plant growth under abiotic stress conditions such as drought, salinity, metal toxicity, and elevated temperatures was suggested (de Zélicourt et al, 2018; Numan et al, 2018). The interaction between plants and endophytes leads to enhanced plant growth under heat stress conditions (Marquez et al, 2007). However, the molecular mechanisms of induced thermotolerance by beneficial microbes are unknown. Here, we report that an endophytic bacterium, *Enterobacter* sp. SA187, isolated from root nodules of the indigenous desert plant *Indigofera argentea* (Andrés-Barrao et al, 2017), significantly enhanced thermotolerance in the model plant *Arabidopsis thaliana*, demonstrating that SA187 has the potential to improve agriculture under extreme desert conditions. SA187 regulates the transcription dynamics of selected heat memory genes in a process that is similar to heat acclimation by hypermethylating histone H3K4 at *APX2* and *HSP18.2* gene loci. Acclimation induced priming of memory genes such as *APX2* and *HSP18.2* depends on the transcription factors *HSFA1A, B, D, and E* and the downstream master regulator *HSF2A* (Lamke et al, 2016; Lamke and Baurle 2017). *hsfa1a,b,d,e* and *hsfa2* mutants revealed that both heat acclimation and bacterial priming function via the same pathway of *HSF* transcription factors to induce heat tolerance in *A. thaliana*. In summary, our study highlights the mechanism of thermotolerance provided by the beneficial plant endophyte SA187 in plants.

## Results

### *Enterobacter* sp. SA187 mediates heat stress tolerance in *Arabidopsis thaliana*

To investigate the impact of SA187 colonization and heat stress acclimation (ACC) on protecting *Arabidopsis thaliana* plants to heat stress, we evaluated fresh weight, % survival, % bleaching and green leaves after heat treatments as our readouts for thermotolerance. Briefly, after five days of germination, SA187- and mock-inoculated seedlings were transferred to ½ MS agar plates. We compared non-heat stressed (NHS) to acclimated (ACC) plants upon direct heat stress of 44°C (HS) with and without bacteria (+/- SA187). For ACC treatment, 9 days old plants that were grown at 22°C were treated at 37°C for 3 h. After two days of recovery at 22°C, a 44°C heat stress was applied for 30 minutes on day 11. After incubation of four days at 22°C, plants were analyzed for phenotypes and transcriptomes. For HS treatment, 11 day old plants that were grown at 22°C, were directly exposed to 44°C for 30 min before further incubation for four days at 22°C and inspected on day 15. Non heat stressed (NHS) plants were grown at 22°C for 15 days in parallel as control reference (Fig 1 A-B). At day 15, we quantified fresh weight, % bleached/green leaves and survival of plants upon the different heat treatments.

**Figure 1:**
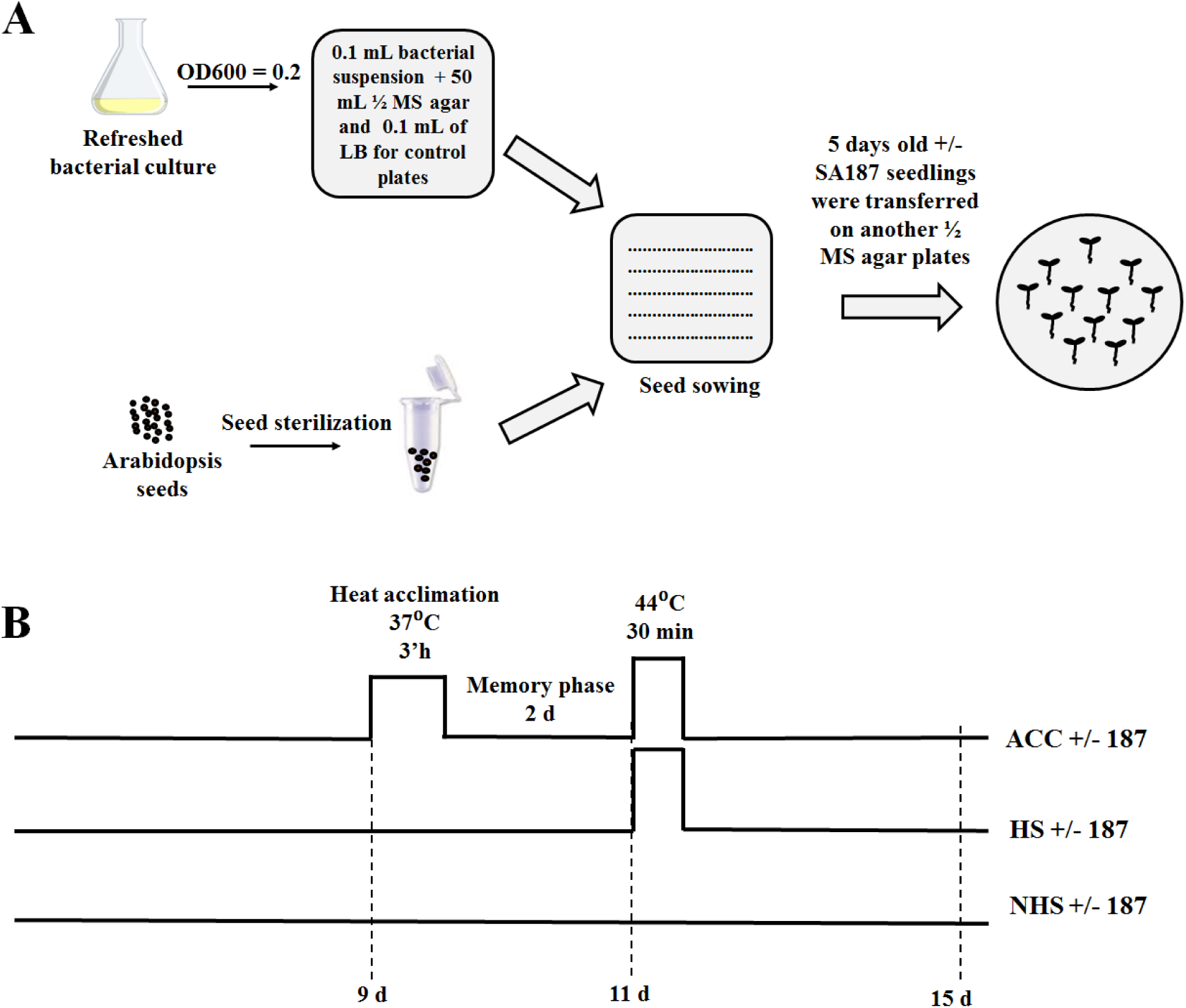
(A) Experimental scheme of heat experiments. 5 day old seedlings +/-SA187 were transferred to new ½ MS plates before (B) heat stress treatment. Heat stress acclimation (ACC): 9 days old plants that were grown at 22°C were treated at 37°C for 3 h, returned to 22°C for 2 days. At day 11, plants were heat stressed at 44°C for 30 min and incubated for 4 days at 22°C before inspection at day 15. Direct heat stress (HS): 11 d old plants that were grown at 22°C were treated at 44°C (HS) for 30 min and incubated for 4 days at 22°C before inspection at day 15. No heat stress (NHS): control plants were grown in parallel at 22°C for 15 d before inspection.

Compared to NHS plants, HS treatment resulted in a major reduction of plant fresh weight and survival as seen in the number of bleached leaves and the ACC treatment significantly protected plants from HS (Fig 2A-E). Comparing SA187-colonized (HS+187) to non-colonized plants (HS), HS+187 exhibited 56.71% higher fresh weight, 25.91% better survival and 34% more of green leaves (Fig 2 A-D). This effect is HS specific, as under control conditions of 22°C, SA187-colonized (NHS+187) and non-colonized plants (NHS) displayed comparable growth, fresh weight and survival levels (Fig 2 A-E). SA187-colonized plants that were in addition heat acclimated (ACC+187) showed the highest fresh weight, survival and green/bleached levels among all the heat stress treatments (Fig 2A-E). These data show that SA187 protects *Arabidopsis thaliana* from heat stress.

**Figure 2:**
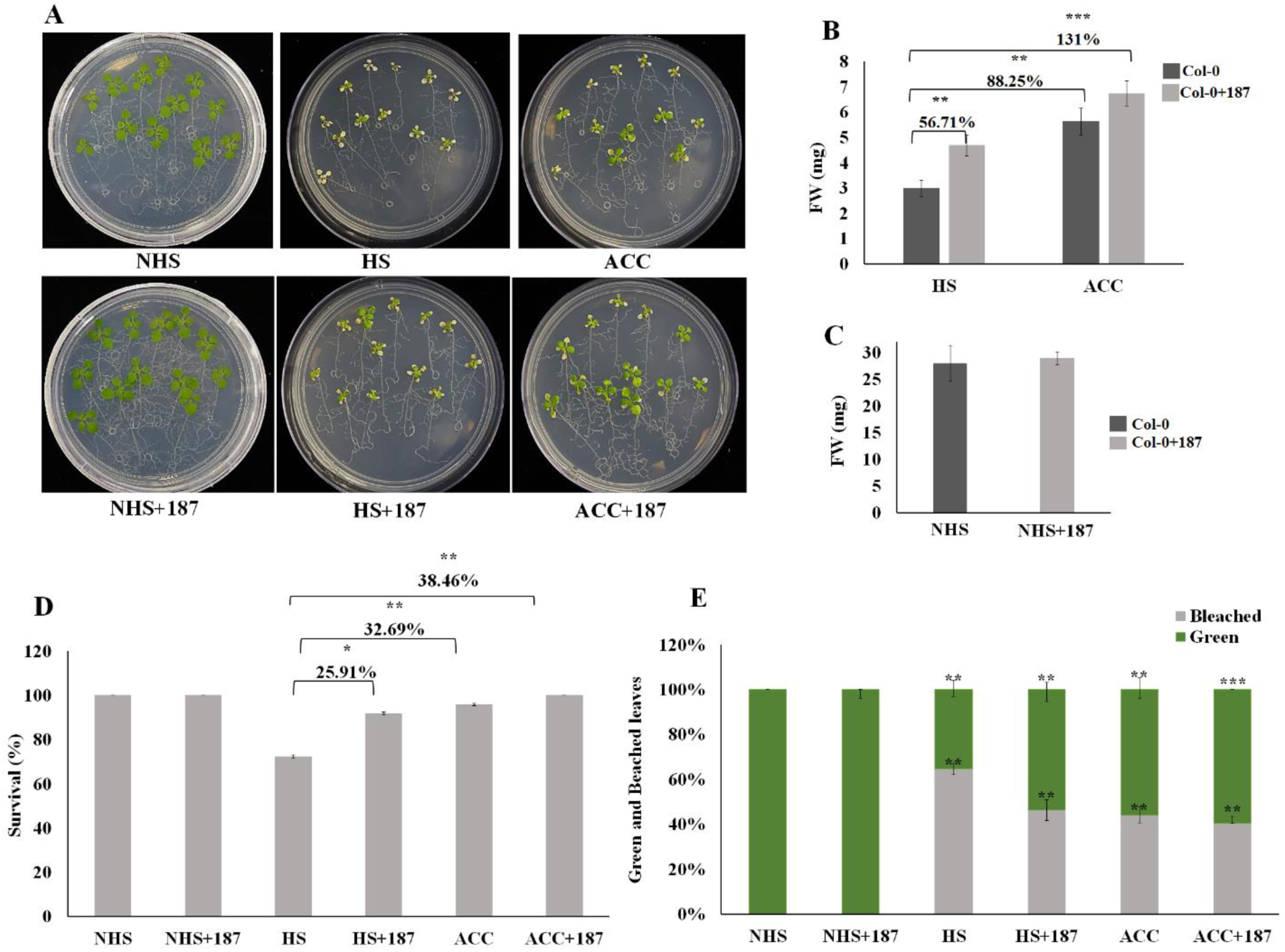
*Enterobacter* sp. SA187 increases thermotolerans in *Arabidopsis thaliana*. (A) The phenotype of *Arabidopsis* seedlings with and without SA187 under different temperature regimes. Left, plants under the normal conditions (22°C) with and without bacteria (NHS, NHS+SA187). Middle, plants without SA187 underwent to a direct heat shock of 44°C (top) and with SA187 (bottom). Left, plants underwent to acclimation treatment with and without SA187 (ACC, ACC+SA187). (B) Fresh weight quantification of plants under two different heat stress, at day 15 after 4 days of recovery. (C) Fresh weight under normal condition. (D) Percent survival of plants. (E) Scoring of bleached (top) and green leaves (bottom) (average of 36 plants) in plants with and without SA187 under two HS and ACC treatments, all the treatments are compared with direct 44°C heat stress treatment. All plots represent the mean of 3 biological replicates (n=36). Error bars represent SE. Asterisks indicate a statistical difference based on the Student’s t-test (* P < 0.05; ** P < 0.01; *** P < 0.001).

### Overview of the transcriptional response of *A. thaliana* during heat stress

In order to determine the genome-wide extent of heat stress-induced changes, we performed transcriptome profiling of SA187-colonized and non-colonized 15 day old plants under non-heat stress (NHS), heat acclimated (ACC) and direct heat stress (HS) conditions (Fig 1B). In comparison to NHS plants, and considering a log2FC ≥ 2 or ≤ - 2 (P < 0.05), HS displayed 4609, ACC 1704, HS+187 2130, while ACC+187 1740 DEGs (Figure 3 A-B, Appendix Table S1). Among all the treatments, HS treatment showed a maximum number of 2627 DEGs that were unique to 44°C treatment. We performed gene enrichment analysis using the AgriGO platform for *A. thaliana*. The gene ontology (GO) terms (FDR≤ 0.05) for 1436 up-regulated, HS DEGs indicated enrichment in single organism process, response to stress, organic substances, signal transduction, protein phosphorylation, response to abscisic acid, temperature, water and cell death. While the GO enrichment for the 1191 down-regulated HS DEGs showed enrichment for cellular, metabolic, phosphorus metabolic, and oxidation-reduction process, photosynthesis, ion transport, growth, response to auxin and gibberellin (Fig. 3 A-B).). The gene enrichment analysis for the common genes in all heat stress treatment is presented in Fig EV1.

**Figure 3:**
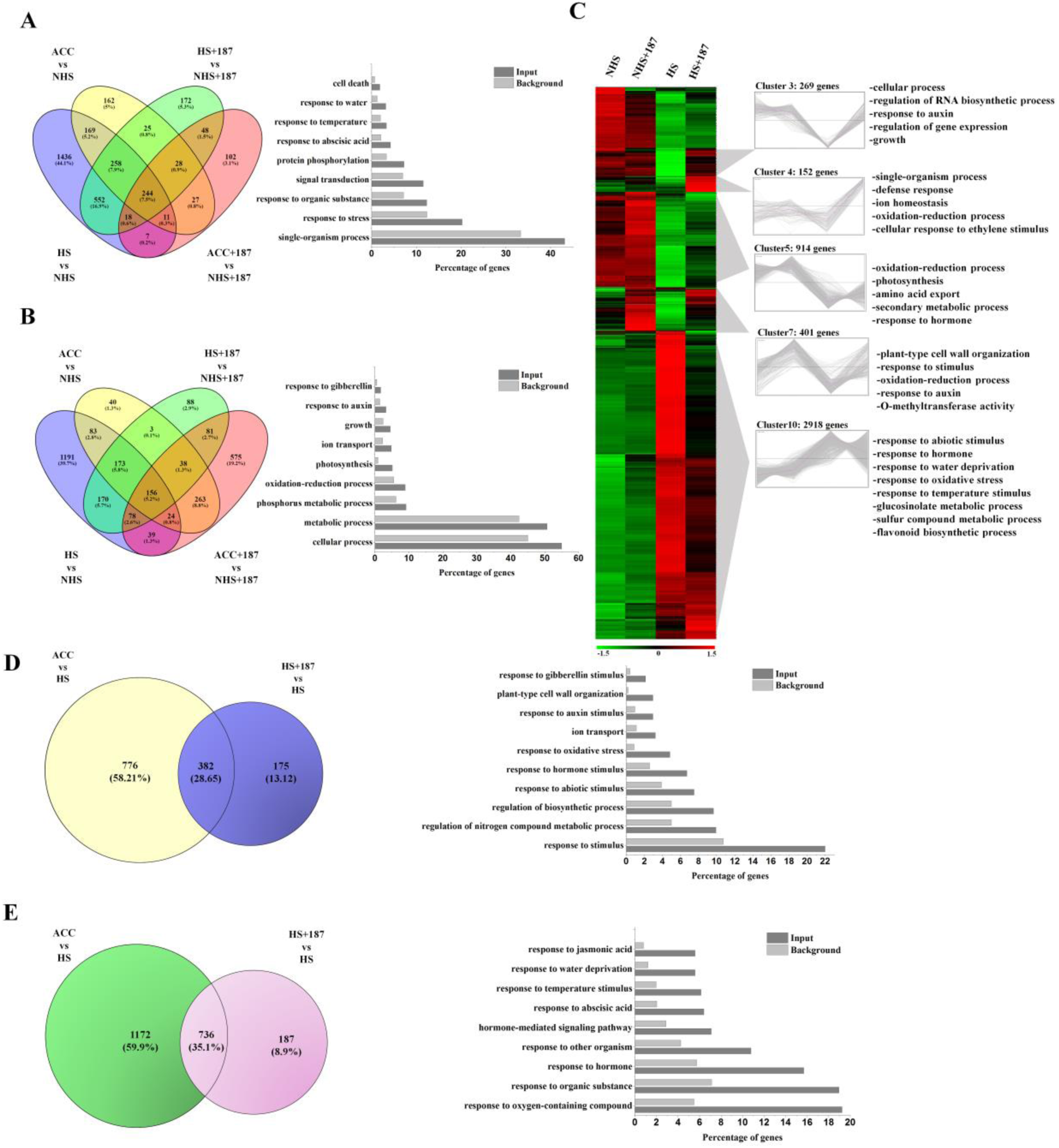
Transcriptome analysis of Arabidopsis response to SA187 under heat stress conditions. (A-B) Venn diagrams representing the number of DE up- and down-regulated genes in response to heat stress (HS, ACC) with and without SA187 compared to NHS and NHS+187. The histograms showing enriched GO terms for unique DE up- and down-regulated genes in HS compare to NHS. (C) Hierarchical clustering of up- and down-regulated genes in *Arabidopsis* seedlings in response to HS and HS+187 treatments based on the RNA-Seq analysis. For every gene, FPKM values were normalized. Red bars denote an increase in expression while green bars indicate a decrease in expression for a given gene. For the most relevant clusters, gene families significantly enriched are indicated based on gene ontology. (D-E) Venn diagram showing the number of DEGs commonly up-regulated and down-regulated in response to HS+187 and ACC compared to HS. The histograms represent the enriched GO terms associated with the DEGs.

### *Enterobacter* sp. SA187-colonized plants respond differently to HS

Next, we aimed to investigate how *Enterobacter* sp. SA187 affects the transcriptome of *A. thaliana* upon heat stress. Under non heat stress conditions, only 303 genes were differentially expressed in SA187-colonized plants (Appendix Table S2). Compared to ambient control conditions (NHS), 4609 genes were found to be differentially expressed upon heat stress (HS) in non-colonized plants, while only 2130 DEGs were found in SA187-colonized plants under the same conditions (HS+187) (Appendix Table S2).

To elucidate the specific role of SA187 in heat stress, the transcriptome data were organized by hierarchical clustering into 10 groups and analyzed for gene ontology enrichment (Fig 3 C). Cluster 3 contains genes that are down-regulated under HS but not in SA187-colonized plants (HS+187) showing enrichment in cellular process, regulation of RNA biosynthesis, response to auxin and regulation of gene expression. Interestingly, cluster 4 genes are significantly induced by SA187 specifically under heat stress conditions. The GO analysis of these 152 genes showed a potential involvement in single-organism process, defense response, ion homeostasis, oxidation-reduction processes and cellular response to ethylene. In addition, a specific effect of SA187 on the transcriptome of plants was found in cluster 7 which consists of 401 DEGs representing genes that are up-regulated by SA187 independently of the growth conditions. This cluster is significantly enriched for GO terms of plant-type cell wall organization, response to stimulus, oxidation-reduction processes, response to auxin and O-methyltransferase activity. The DEGs of cluster 10 are slightly affected by SA187, by having lower fold change expression levels. Cluster 5 with a number of 914 DEGs displayed down-regulation of genes in HS and HS+187, and are mainly involved in the oxidation-reduction process, photosynthesis, amino acid export, secondary metabolism and response to hormone. Finally, cluster 10 comprised the largest set of differentially expressed genes that are strongly up-regulated by HS but show only moderate induction by HS+187. These genes mainly consist of heat stress regulated genes which show enrichment for response to abiotic stimulus, hormone, water deprivation, oxidative stress, temperature stimulus, glucosinolate and sulfur compound metabolism and flavonoid biosynthesis.

We also analyzed the effects of acclimatization by comparing ACC with HS. Of the 3064 DEGs (1157 up- and 1907 down-regulated genes) (Appendix Table S2), the GO terms for up-regulated genes were enriched for regulation of biosynthetic processes, response to auxin, photosynthesis, and response to gibberellin (Fig EV2).

Since our results suggested that SA187 and ACC modulated the *A. thaliana* heat stress transcriptome, we searched for common features between the two treatments by comparing HS+187 vs HS and ACC vs HS. This analysis revealed a strong overlap of 1118 genes between the two treatments (HS+187 vs HS and ACC vs HS) with 382 up- and 736 down-regulated genes (Appendix Table S3). The commonly up-regulated genes showed GO enrichment for the regulation of biosynthesis, cell wall organization, response to the hormone, auxin and gibberellin stimulus, while the down-regulated genes showed GO enrichment categories such as response to stress, temperature stimulus, response to ABA, water, SA and jasmonic acid biosynthesis (Fig 3 D-E). This vast overlap between the two treatments suggested that SA187 and ACC might use common mechanisms to protect Arabidopsis from extreme temperatures.

### *Enterobacter* sp. SA187 enhances the expression of heat-responsive and memory genes upon heat stress

To understand the molecular mechanism by which SA187 confers heat stress resistance to *A. thaliana*, we investigated the expression pattern of 8 heat-responsive genes (*HSP101, HSP70, HSP70b, GA3OX1, ATERDJ3A, HSP90, XTR6* and *MIPS2*) and 3 heat stress memory genes including *HSFA2* which is a master regulator of heat stress and *HSP18.2* and *APX2* which are known to play a vital role in heat stress memory (Liu et al, 2018). We collected samples after 1 h, 24 h and 48 h of exposure to 44°C and their respective controls at 22°C (Fig 4 A). Under control conditions, SA187 (NHS+187) did not change the expression of these heat responsive and memory genes in comparison to non-colonized plants (NHS) (Fig 4 B and C). On the contrary, after 1 h of 44°C (HS), SA187-colonized (HS+187) and ACC plants showed significantly higher levels of transcripts for all heat responsive genes than HS treated plants (both non-colonized and non-acclimated plants) (Fig 4 B). However, at 24 h after the 44°C treatment, the transcript levels in SA187-colonized plants (HS+187) were either lower (*HSP101, ATERDJ3A, HSP90*) or similar to HS (*HSP70, HSP70b, XTR6, MIPS2*). At 48 h of HS, the transcript levels stayed similar for both HS+187 and HS, but declined relative to levels at 1 h and 24 h of heat stress. Similarly, for ACC treated plants, at 24 and 48 h, *GA3OX1, XTR6* and *MIPS2* transcript levels were higher than in HS plants, while for the other 5 genes (*HSP101, HSP70, HSP70b, ATERDJ3A, HSP90*) we observed lower expression values compared to HS. Likewise, for 3 heat stress memory genes, compared to HS plants, SA187-colonized (HS+187) and heat acclimated (ACC) plants exhibited significantly higher levels of transcripts for *HSFA2, HSP18.2* and *APX2*. However, at 24 and 48 h of heat stress, the expression levels for *HSFA2* had declined when compared to HS, but surprisingly not for *HSP18.2* and *AXP2*, which stayed significantly higher until 48 h after heat stress (Fig 4 C). This pattern of gene expression possibly explains why the SA187-colonized and heat acclimated plants showed better survival to high temperatures than control plants.

**Figure 4:**
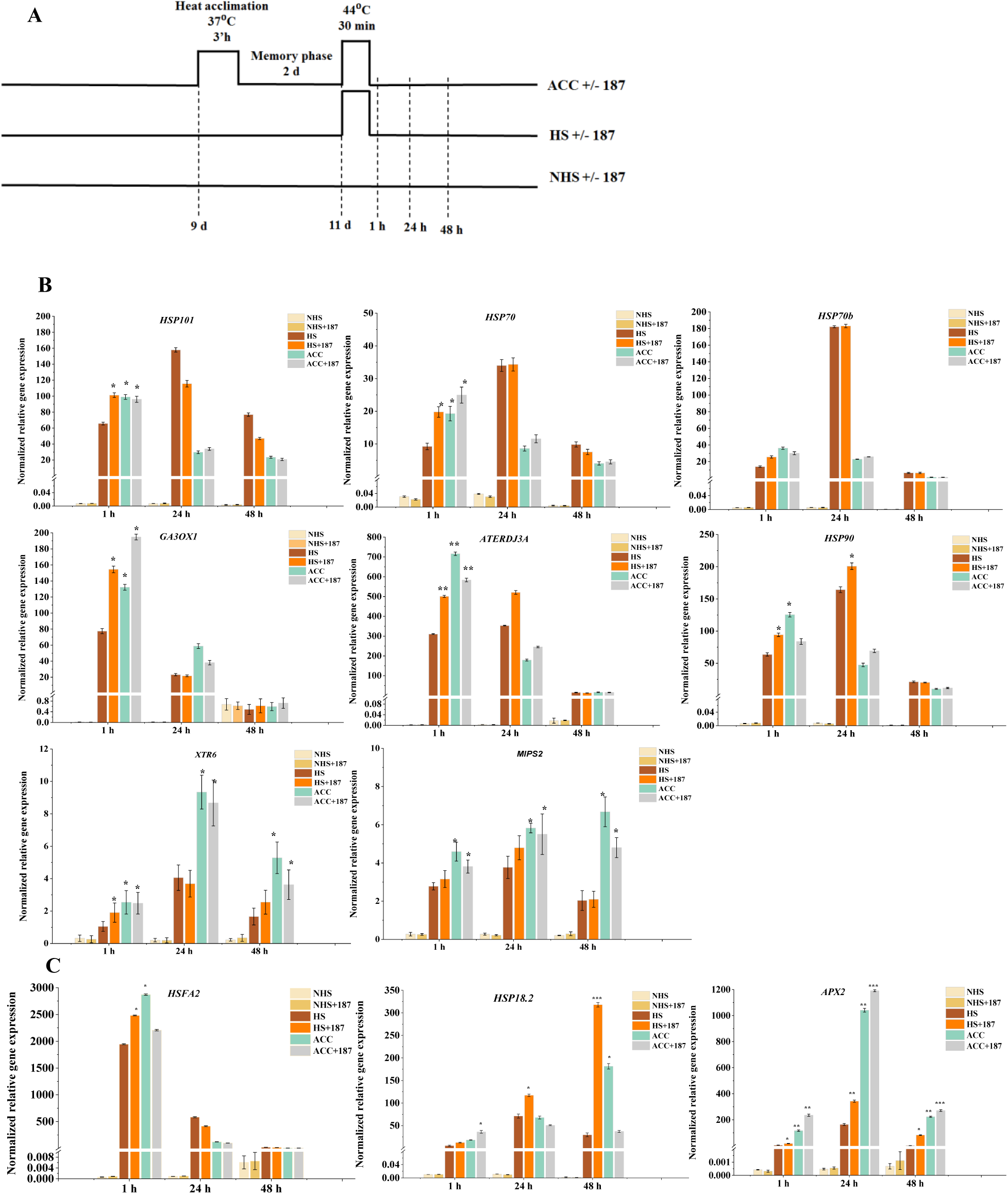
*Enterobacter* sp. SA187 up-regulates the expression of heat-responsive and memory genes upon direct exposure to 44°C heat stress. (A) A graphic presentation of different treatments and sampling time points selected for qRT-PCR. (B) Heat-shock proteins (*HSP*s), and other heat responsive genes show higher transcript levels by presence of SA187 (HS+187) and acclimation (ACC) in comparison to plants exposed to direct 44°C heat stress (HS) after 1 h of HS, while at 24 and 48 h all these genes showed equal or lower transcript levels than non-inoculated or non-acclimated plants. (C) Transcript levels of *HSFA2, HSP18.1* and *APX2* in control (NHS, NHS+187), HS+187, ACC and ACC+187 treated plants at 1, 24 and 48 h of direct HS. Transcript levels were normalized to reference gene tubulin and the respective 22°C NHS harvested at the same time point. All the treatments are compared with direct 44°C heat stress treatment for statistical significance. All plots represent the mean of 3 biological replicates. Error bars represent SE (* P≤ 0.05 and ** P≤ 0.01).

### *Enterobacter* sp. SA187 induces sustained H3K4me3 levels at *Arabidopsis APX2* and *HSP18.2* gene loci

SA187-colonized and heat-acclimated plants showed prolonged higher levels of *APX2* and *HSP18.2* transcripts than non-colonized or non-acclimated plants (Fig 4 C). Recently, chromatin modifications were shown to be involved in maintaining the prolonged expression of heat-responsive memory genes, such as *APX2* and *HSP18.2* in *Arabidopsis* (Lamke et al, 2016a). In particular, histone H3K4me3 levels were elevated for 3-5 days in these genes after HS acclimation, and this chromatin modification primes plants to tolerate subsequent severe HS (Liu et al, 2015; Lamke et al, 2016a, b). To test whether H3K4me3 is also involved in providing an elevated and prolonged-expression of the heat memory genes in SA187-colonized plants, we evaluated H3K4me3 levels at the *APX2* and *HSP18.2* gene loci as representatives of the HS memory-related genes in 9 and 12 days old plants. We performed ChIP-qPCR of plants that were primed at 37°C for 3 h (P, P+187) and their respective non-primed controls (NP, NP+187) before incubation for 24 and 72 h at 22°C (Fig 5 A-B). As shown in previous studies (Lamke et al, 2016a; Liu et al, 2018), the 37°C primed plants showed enrichment of H3K4me3 at regions 2 and 3 of *APX2* and region 2 of *HSP18.2* gene (Fig 5 C). For *APX2*, H3K4me3 levels remained high until 72 h after 37°C heat stress priming, and for *HSP18.2*, the levels were still elevated, but lower than at 24 h (Fig 5 C). Interestingly, the ChIP assay of SA187-colonized plants (NHS+187) also showed significant enrichment for H3K4me3 at the *APX2* and *HSP18.2* loci (Fig 5 C). When SA187-colonized plants were primed at 37°C (P+187), the H3K4me3 enrichment for *APX2* was higher than in SA187-colonized non-primed plants (NP+187) but less than in 37°C primed (P) plants (Fig 5 C). In summary, SA187 colonization causes H3K4me3 accumulation at HS memory-related gene loci.

**Figure 5:**
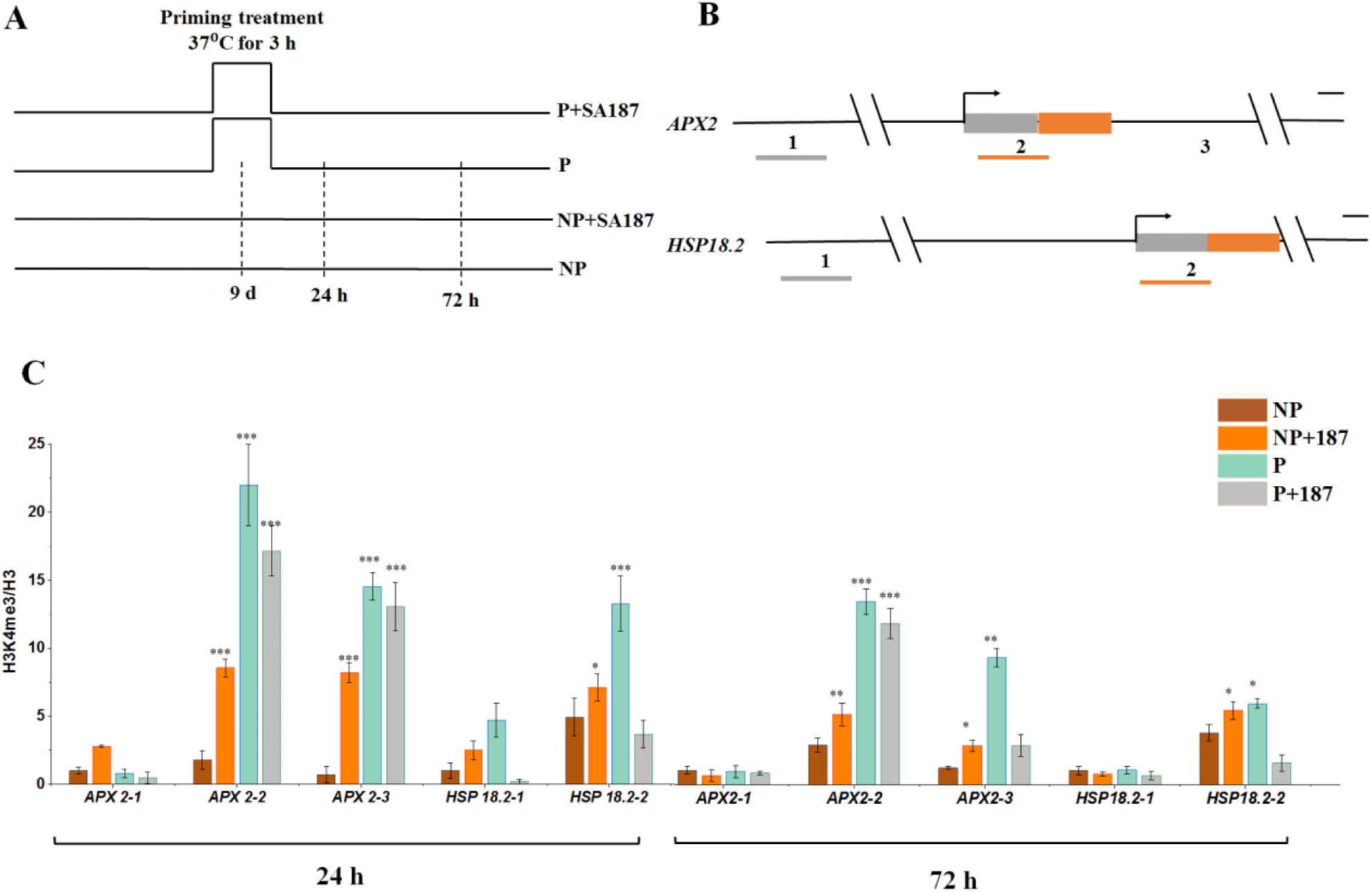
*Enterobacter* sp. SA187-induced heat stress memory gene expression is associated with changes in chromatin modification. (A) A schematic representation of the experimental set-up and sampling times. 9 day old plants were primed by heat treatment at 37°C for 3 h before incubation at 22°C for 24 h or 72 h. (B) *APX2* and *HSP18.2* gene models drawn to scale (grey boxes, 5’ untranslated region; orange boxes, exons; angled arrow, transcription start site). (C) Relative enrichment of H3K4me3 at *APX2* and *HSP18.2* in SA187-colonized non-primed plants (NP+187), 37°C-primed (P) and 37°C-primed SA187-colonized plants (P+187) at 24 and 72 h after priming as determined by chromatin immunoprecipitation-qPCR for the indicated regions of *APX2* and *HSP18.2*. Amplification values were normalized to input and H3 and region 1 of non-primed (NP) plants. *P < 0.05; **P < 0.01; **P < 0.001 for differences between NP in comparison to NP+187, P and P+187 treatments.

### SA187-induced heat stress tolerance is mediated by the master regulator *HSFA2*

Since *HSFA2* is the master regulator of heat stress responsive genes and heat acclimation (Liu and Charng 2012), we wanted to understand whether it also plays a role in SA187-induced heat stress tolerance. As expected, *hsfa2* mutant plants were strongly compromised in heat acclimation (ACC), but the beneficial effect of SA187 was equally compromised, indicating that *HSFA2* is also important for mediating SA187-induced thermotolerance (Fig 6 A-C, Fig EV3 for *hsfa2* control plants). We further investigated the expression of the heat stress-responsive and memory representative genes *APX2, GA3OX1, HSFA2* and *HSP70* in wild type and *hsfa2* mutant plants. The expression of these four genes by HS was tremendously reduced in *hsfa2* mutants as compared to wild type (°). Moreover, in contrast to wild type Col-0 plants, none of the genes showed induction in ACC or SA187-colonized *hsfa2* mutant plants (Fig 7). *HSFA2* expression itself is regulated by the redundantly acting upstream transcription factors *HSFA1A, HSFA1B, HSFA1D* and *HSFA1E* (Liu et al, 2018) and quadruple knockout *hsfa1-q* (*hsfa1a,b,d,e*) mutant plants are hypersensitive to heat stress and unable to induce *HSFA2* dependent gene expression of *APX2, GA3OX1, HSFA2* and *HSP70* upon ACC treatment. To test whether the *HSFA1* transcription factors are also involved in mediating SA187-induced heat stress tolerance, we tested gene expression of *APX2, GA3OX1, HSFA2* and *HSP70* in *hsfa1-q* mutant plants. Similar to ACC treatment, SA187-colonized *hsfa1q* mutant plants were compromised in the induction of *APX2, GA3OX1, HSFA2* and *HSP70* gene expression (Fig 7), further confirming that both ACC- and SA187-mediated thermotolerance are regulated by the same transcriptional network of heat stress transcription factors.

**Figure 6:**
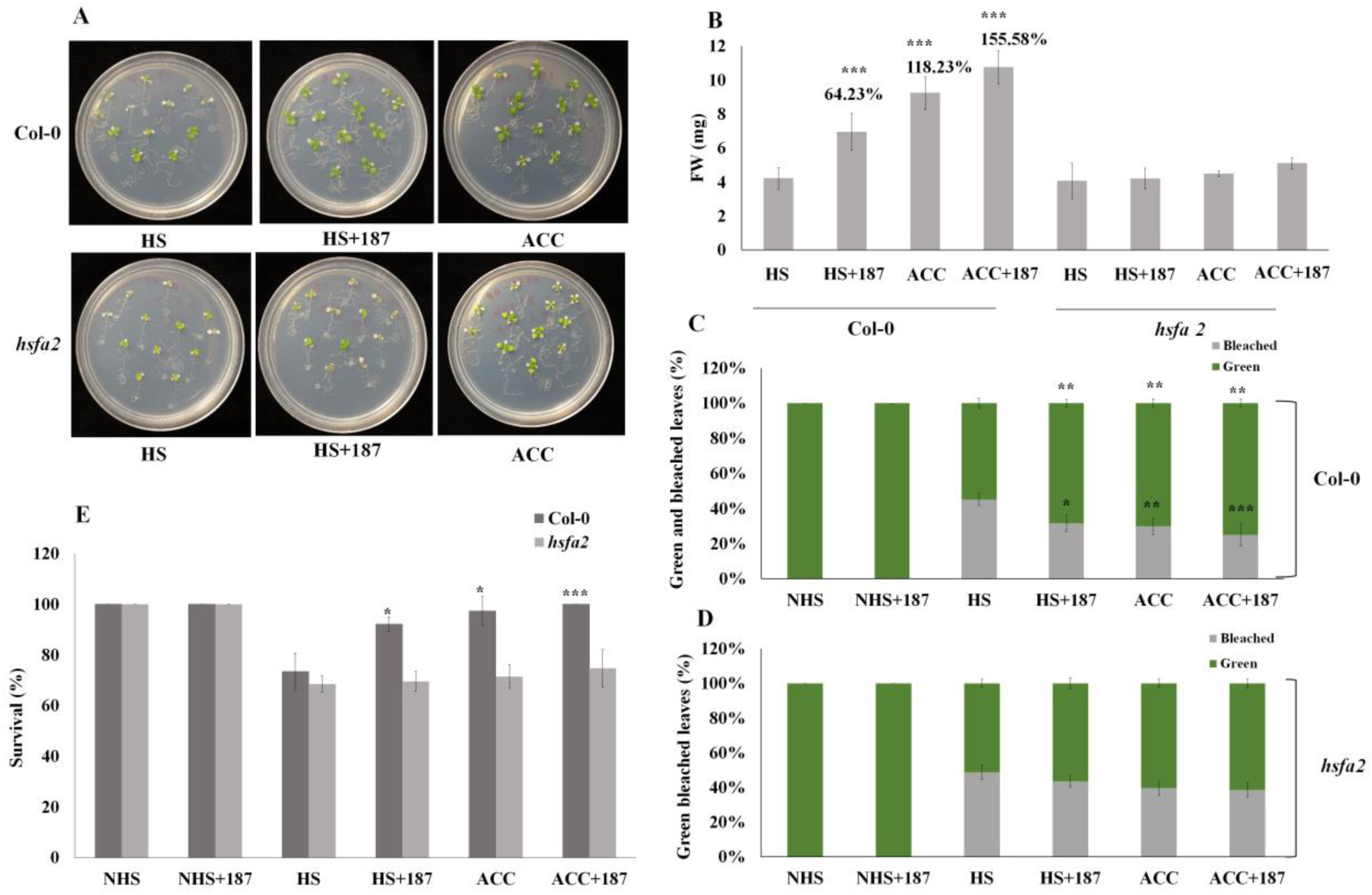
*HSFA2* is essential for the beneficial response of SA187 and acclimation in *Arabidopsis*. (A) The phenotypes of SA187-colonized or non-colonized wild type and *hsfa2* mutant plants upon heat acclimation (ACC) or direct heat stress (HS). HS: Non-colonized 11 day old plants were treated at 44°C for 30 min before incubation for 3 d at 22°C. HS+187: 11 d old SA187-colonized plants were treated at 44°C for 30 min before incubation for 3 d at 22°C. ACC: 9 d old non-colonized plants were primed for 3 h at 37°C before incubation for 2 days at 22°C and heat treatment at 44°C for 30 min before further incubation at 22°C for 3 days. (B) Fresh weight quantification, (C-D) Bleaching (top) and green leaves (bottom) quantification (average of 36 plants) and (E) Percent survival in NHS, NHS+187, HS, HS+187, ACC and ACC+187 plants. All the treatments are compared with HS treatment. All plots represent the mean of 3 biological replicates (n=36). Error bars represent SE. Asterisks indicate a statistical difference based on the Student’s t-test (* P < 0.05; ** P < 0.01; *** P < 0.001).

**Figure 7:**
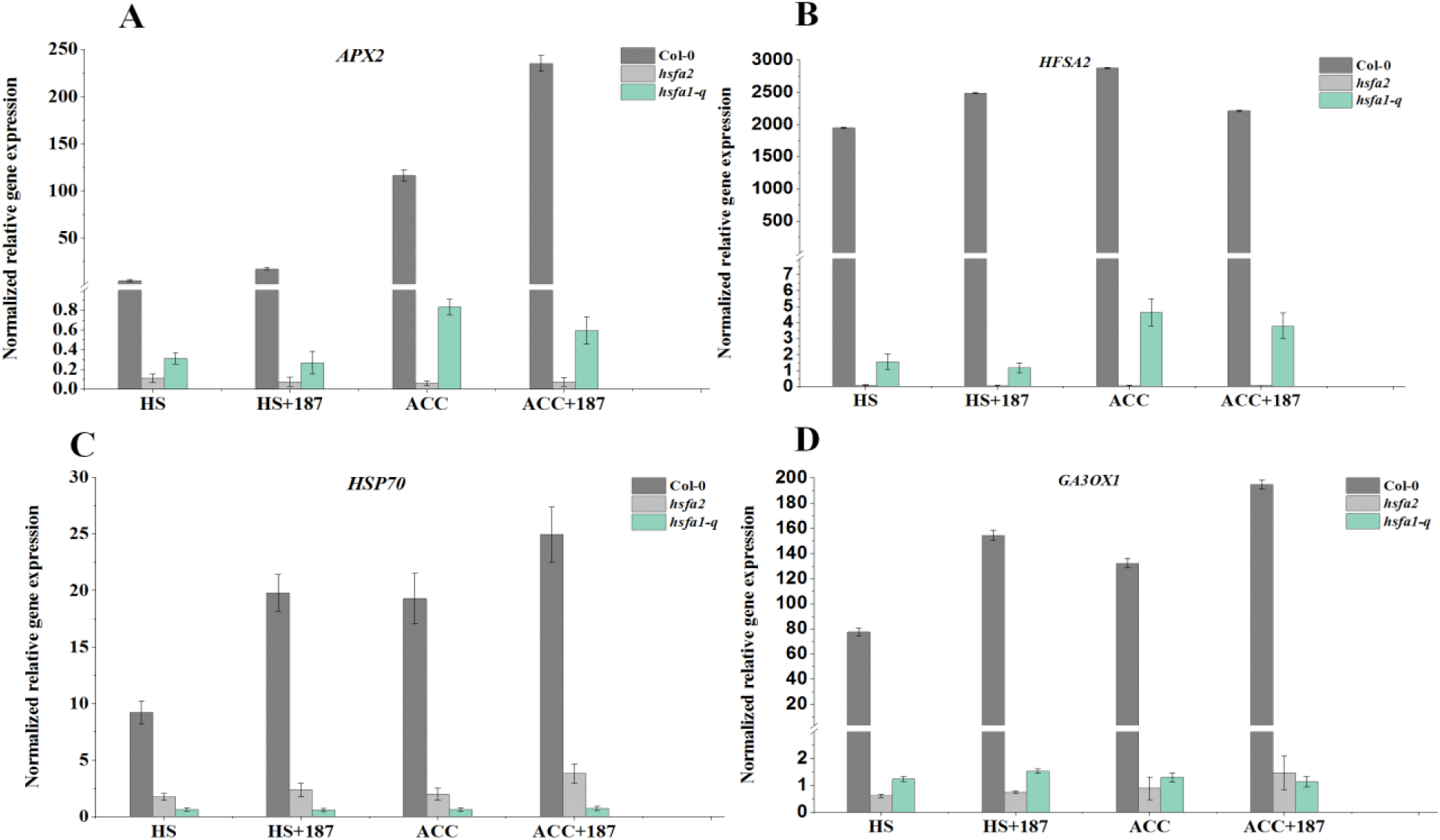
The SA187 and acclimation mediated transcriptional memory after HS depends on functional *HSFA2* and *HSFA1A,B,D,E* (*HSFA1-q)*. Transcript levels of *APX2, GA3OX1, HSFA2* and *HSP70* in Col-0, *hsfa2* and *hsfa1a,b,d,e (hsfa1-q)* mutant plants after 1 h of direct heat stress in HS, HS+187, ACC and ACC+187 treatments. Non-colonized 11 day old plants were treated at 44°C for 30 min before incubation for 3 d at 22°C. HS+187: 11 d old SA187-colonized plants were treated at 44°C for 30 min before incubation for 3 d at 22°C. ACC: 9 d old non-colonized plants were primed for 3 h at 37°C before incubation for 2 days at 22°C and heat treatment at 44°C for 30 min before further incubation at 22°C for 3 days. Transcript levels were normalized to reference gene tubulin and the respective 22°C NHS harvested at the same time point. Error bars represent SE.

## Discussion

The increases in temperature due to global warming have a significant negative impact on agriculture worldwide, and pose a serious threat to global food security. To cope with this challenge, several solutions are proposed, mainly by breeding and genetic modifications of crops for enhancing heat tolerance. However, breeding programs are time-consuming and costly, while gene transformation technology is not well accepted by society (Asseng et al, 2015; Craita and Tom 2013; Ling et al, 2018). Hence, beneficial microbes might be an interesting alternative to enhance heat tolerance of plants. Several studies reported that endophytes can enhance plant growth under heat stress conditions (Marquez et al, 2007, de Zelicourt, et al, 2013; 2018, Eida et al, 2018). So far, however, the molecular mechanisms underpinning the acquisition and maintenance of plant heat stress by beneficial microbes were not investigated. To understand these molecular mechanisms, we studied *Enterobacter* sp. SA187-induced heat stress tolerance in the model plant *A. thaliana*. In the present study, we demonstrated that SA187 can significantly increase heat resistance in *Arabidopsis* plants. SA187 can interact in a beneficial way with other non-host plants, such as *Medicago sativa*, in open field desert agriculture (de Zelicourt et al, 2017), suggesting that beneficial microbes such as SA187 might be efficient means to improve global crop productivity at higher temperatures.

To understand the interaction between SA187 and *A. thaliana* under heat stress conditions at the molecular level, we performed RNA-seq analysis of plants that were incubated for 3 days at 22°C after a severe heat stress treatment of 22°C for 30 min. Although both ACC and SA187-colonized plants showed some overlap with the HS gene set of naive NHS plants, the majority of the genes induced by HS were not induced in SA187-colonized and acclimated plants (ACC vs NHS and HS+187 vs NHS, Fig 3 A and B). The HS-specific gene sets in Fig 3 A and B showed GO enrichment of up-regulated genes for response to stress, organic substances, signal transduction, protein phosphorylation, response to abscisic acid, temperature, water and cell death (Fig 3 A), while the down-regulated genes showed enrichment for metabolic, phosphorus metabolic and oxidation-reduction process, photosynthesis, ion transport, growth, response to auxin and gibberellin (Fig 3B). These features correspond to the phenotype of HS treated plants showing growth arrest and enhanced cell death (Fig 2). In contrast, ACC and SA187-colonized plants did not show differential expression of these genes and resumed growth at 4 days after severe heat stress (Fig 2), suggesting that the exclusive expression of the HS-specific gene set in HS plants might be associated with the enhanced cell death in NHS and non-colonized plants.

The comparison of the transcriptomes of SA187-colonized to non-colonized, plants the transcriptomes of heat acclimated (ACC) plants to those of non-acclimated (HS) plants 4 days after the severe heat stress should reveal the specific contribution of SA187 to heat stress tolerance. These comparisons revealed that the genes associated with SA187-induced heat stress tolerance largely overlapped with those of ACC plants (Fig 3 D and E), strongly indicating that SA187- and ACC-induced heat stress tolerance should use similar gene networks for heat protection. Interestingly, the GO terms of the common segments of the two processes are associated with the upregulation of genes associated with the hormones auxin and gibberellin, while those of the down-regulated common genes are linked to jasmonic and abscisic acid. The function of these hormones in heat stress tolerance in general and in particular in ACC- and SA187-induced protection is not clear yet and deserves further attention.

In order to understand the mechanisms that are responsible for SA187-induced plant survival under heat stress conditions, we also performed a targeted transcriptome analysis for some of the heat-responsive and memory genes. For ACC, it has been shown that acquired thermotolerance is associated with a transcriptional memory resulting in faster and stronger expression of heat-responsive genes upon a repeated stress signal (Stief et al, 2014; Avramova, 2015; D’Urso and Brickner, 2017). It is believed that the transcriptional memory of past exposure to environmental stress may serve the plants to be better prepared for a future stress incident. In the present study, we show that SA187-induces heat stress tolerance to a similar degree as ACC and in both cases, hyper-induction of heat-responsive and heat memory genes is observed upon a severe heat stress challenge. The hyper-induction of heat-responsive genes is proven to be responsible for the better performance of ACC plants (Lamke et al, 2016a; Liu et al, 2018; Ling et al, 2018). Moreover, it was shown that *HSFA2* is required for the maintenance of heat stress-induced memory by activating the expression of HS genes (Wu et al, 1995; Schramm et al, 2006; Lamke et al, 2016a, 2016b). Interestingly, we also found that *HSFA2* and other heat responsive genes (*HSP101, HSP70, 70b, GA3OX1, HSP90.1*, and *ATERDJ3A*) were induced by the presence of SA187 bacteria at 1 h after HS. However, at 24 and 48 h, non-colonized or non-acclimated plants showed higher expression for some of the genes, indicating that in heat- and bacteria-primed plants gene expression patterns were reset back to normal levels once the stress was relieved except for non-primed plants. Moreover, in both SA187-colonized and heat-acclimated plants, gene expression of the heat memory genes *APX2* and *HSP18.2*, which are known to be involved for maintaining heat stress memory (Lamke et al, 2016a; Sedaghatmehr et al, 2016), were up-regulated and higher transcript levels were maintained in SA187-colonized and heat-acclimated plants up to 48 h. These data suggest that different genes are involved in rescuing plants from heat stress and that the transcriptional memory depends on prolonged higher transcript levels of some specific gene. In summary, the gene expression data suggest that SA187 elevates the transcript levels of heat memory-related genes thereby helping plants to survive better to acute heat stress.

The transcriptional regulation of heat responsive genes is known to be controlled by epigenetic factors that help maintain priming memory (Berry and Dean, 2015). Such patterns must be maintained to confer thermotolerance to heat stress. For transcriptional memory several mechanisms have been proposed such as chromatin modification of histones or the use of histone variants (Brickner et al, 2007; Laine et al, 2009; Tan-Wong et al, 2009; Light et al, 2013). A number of reports showed a role of epigenetic regulators in maintaining plant heat stress memory (Liu et al, 2015) and elevated histone H3K4 methylation levels and enhanced chromatin accessibility was shown to be involved in the memory process induced by acclimation (Brzezinka et al, 2016; Lamke et al, 2016a). An involvement of chromatin modifications in providing heat stress tolerance by beneficial microorganisms is hitherto unknown. In this work, we show that similar to heat acclimation priming, the beneficial bacterium *Enterobacter* sp. SA187 also induces sustained accumulation of H3K4me3 at *APX2* and *HSP18.2* gene loci (Fig 5 C). The accumulation of H3K4me3 is particularly high and long-lasting at *APX2* while it is somewhat lower at *HSP18.2* gene loci. Interestingly, this pattern appears to correlate with the transcript levels in SA187-colonized and heat-acclimated plants. *APX2* (ascorbate peroxidase 2) is known for its role in scavenging reactive oxygen species (ROS), and upregulation may be necessary to increase ROS scavenging activity promptly after the onset of heat stress to prevent cellular damage (Liu et al, 2018). These results are in alignment with previous studies that *HSF*s and *HSP*s are required for providing heat resistance to plants and their transcript levels are affected by H3K4me3 (Liu et al, 2018; Lamke et al, 2016a).

*HSFA2* is a major regulator of HS memory, since it is the most strongly heat-induced *HSF* in *A. thaliana*, and its induction depends on functional *HSFA1* isoforms (Liu et al, 2011; Nishizawa-Yokoi et al, 2011; Liu and Charng, 2013). Furthermore, acclimation-induced heat tolerance depends on *HSFA2* as *hsfa2* mutants are severely compromised in acclimation. SA187-induced heat stress tolerance was similarly abrogated in *hsfa2* mutants, indicating that both SA187-induced heat stress tolerance and heat acclimation are mediated by *HSFA2*. Moreover, the expression of *APX2, HSFA2, GA3OX1* and *HSP70* genes in *hsfa2* mutant cannot be induced any more by heat acclimation, indicating the essential role of *HSFA2* in heat stress memory. Importantly, SA187-colonized quadruple *hsfa1-q* mutant plants also lost SA187-induced heat induction of the *APX2, HSFA2, GA3OX1* and *HSP70* genes, indicating that SA187-induced heat tolerance is regulated in a similar manner as heat acclimation via *HSFA1A, B, D* and *E* and the downstream master regulator *HSFA2* (Fig 7). Therefore, we propose a model that heat acclimation and SA187 functions in a similar mode via *HSFA1-q, HSFA2* dependent transcription of heat responsive genes and H3K4me3 mediated longer persistence of heat memory genes (Fig 8). In terms of applying these findings to agriculture, an important difference exists between SA187-induced heat tolerance and heat acclimation. To obtain plant heat tolerance by heat acclimation, plants need to be pretreated at lower temperatures and the heat stress memory usually only lasts for several days. It is therefore a transient mechanism that is hard to apply on crops that grow on a field. In contrast, SA187 permanently colonizes its host, rendering plants constitutively heat stress tolerant without any further treatments, making this a potential tool to meet the challenges of adapting crop production under global warming conditions.

**Figure 8:**
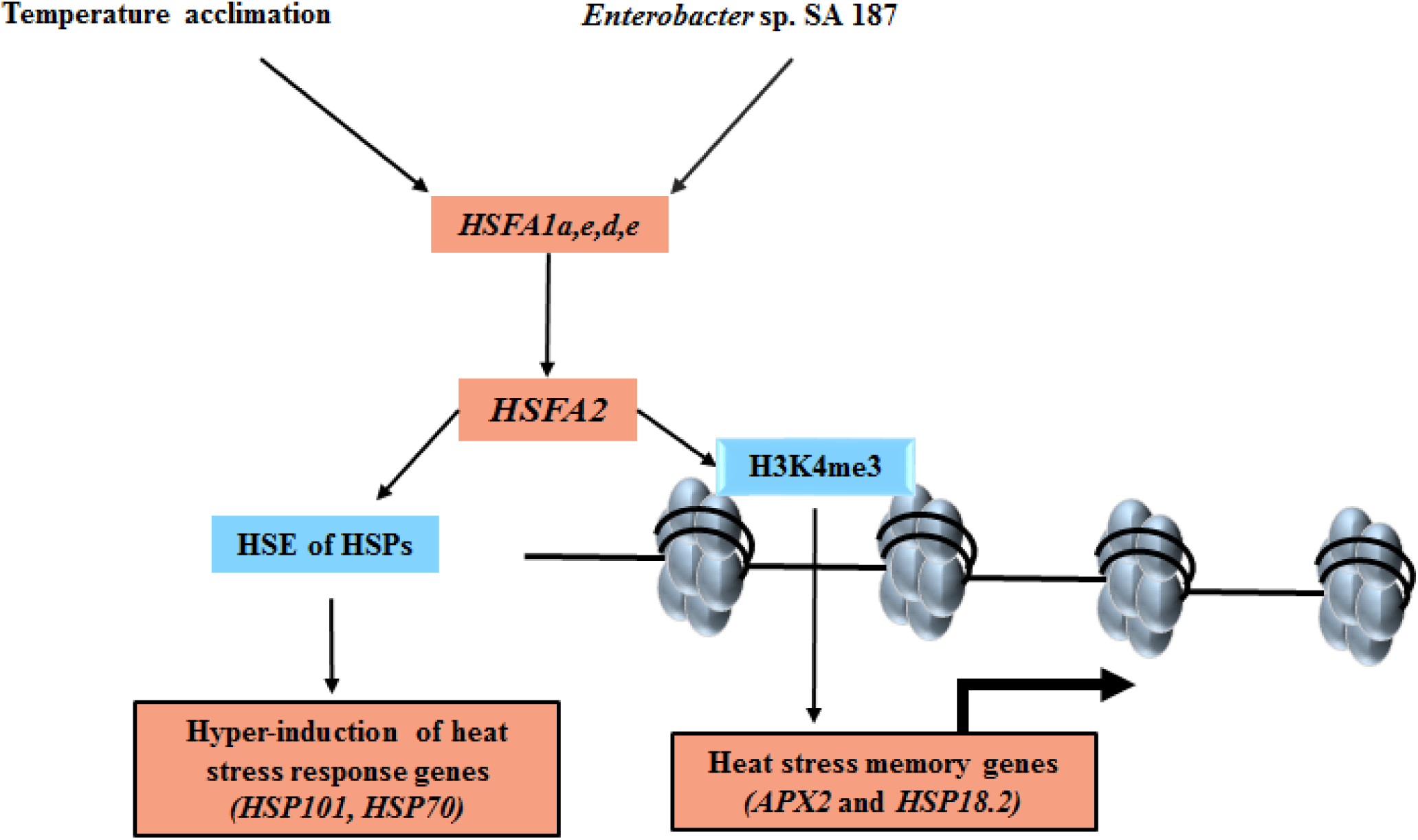
Proposed model of SA187 mediated thermotolerans in *A. thaliana* via *HSFA1, HSFA2* and chromatin modification regulated gene expression of heat-responsive genes.

## Material and methods

### Bacterial inoculum and media preparation

*Enterobacter* sp. SA187 was isolated from root nodules of *Indigofera argentea* in the Jizan region of Saudi Arabia (Andrés-Barrao et al, 2017). Cryogenically maintained *Enterobacter* sp. SA187 were streaked out on LB agar media and incubated at 28°C for 24 h. Streaked bacterial culture was used for further experiments. For bacterial seed plates, 50 mL of half-Murashige and Skoog medium (MS) with 0.9% agar and a pH of 5.8 were mixed with 0.1 mL of fresh bacterial suspension with an OD of 0.2 to obtain a final number of 10^5^ CFU mL^-1^. For control plates, 0.1 mL of liquid LB was mixed with ½ MS media (Fig 1 A).

### Plant material and growth conditions

*Arabidopsis thaliana* Col-0 seeds were obtained from publicly available collections. *hsfa1-q* and *hsfa2* seeds were obtained from Yee-yung Charng (ABRC, Taipei, Taiwan). Seeds were surface-sterilized for 10 min with 0.05% SDS solution prepared in 70% ethanol. The sterilized seeds were plated on ½ MS medium agar plates seeded with SA187 and mock (LB). Seeds were stratified at 4°C for 2 d and then plates were transferred to a growth chamber (Model CU36-L5, Percival Scientific, Perry, IA, USA) under a 16-h photoperiod and 8-h dark conditions at 22°C for germination and seedling growth.

### Heat priming and heat-shock experiment

In the present study we developed a heat-priming platform by modifying previous studies (Larkindale and Vierling, 2008). Five days old SA187-inoculated and non-inoculated seedlings of almost equal lengths were transferred onto new ½ MS plates. For heat stress treatments, plants with bacteria and without bacteria were divided into three sets of plates. In set 1, plants were given acclimation heat stress treatment, where 9 days old SA187-colonized and non-colonized plants were exposed to 37°C of heat acclimation for 3 h followed by 2 d recovery at 22°C and a further 44°C heat stress at day 11 (ACC). In set 2, SA187-inoculated and non-inoculated plants were exposed directly to 44°C heat stress on day 11 (HS). For 44°C treatment, we used water-bath. In set 3, SA187-inoculated and non-inoculated plants were grown under normal conditions at 22°C (NHS). We performed the same heat stress procedure for all experiments and the arrows indicate the sampling points performed for respective data analysis (Fig 1 B).

### RNA extraction, reverse transcription and qRT-PCR

For targeted gene expression study, the plant RNA was extracted from 11, 12 and 13 days old seedlings using the Nucleospin RNA plant kit (Macherey-Nagel), including DNaseI treatment, and following manufacturer’s recommendations. The total RNA was reverse-transcribed using a using SuperscriptIII (Invitrogen): 1 μg of total RNA, oligo-dT as a primer, following manufacturer’s recommendations. For *Arabidopsis* gene expression analyses, tubulin was used as a reference gene. All reactions were done in a CFX96 Touch Real-Time PCR Detection System (BIO-RAD) as follows: 50°C for 2 min, 95°C for 10 min; 40× (95°C for 10 sec and 60°C for 40 sec). All reactions were performed in three biological replicates, and each reaction as a technical triplicate. A list of all primers used in the current study is provided in Appendix Table S4.

### Bioinformatics analysis of RNA-seq data

We performed mRNA libraries with 1 µg of total plant RNA using a stranded mRNA Library Prep kit (Illumina). Pooled libraries were sequenced using Illumina HiSeq 4000 platform which resulted in paired-end reads of length 151 bps. Sequenced reads were checked for quality using FASTQC (Andrews, 2012). Adapter sequences and low-quality reads or base-pairs were trimmed using Trimmomatic V0.36 (Bolger AM et al, 2014). The Parameters for read quality filtering were set as follows: minimum length of 36 bp; Mean Phred quality score greater than 30; Leading and trailing bases removal with base quality below 3; Sliding window of 4:15. Trimmed reads were then aligned to the TAIR10 and SA187 genome combined using TopHat v2.1.1 (Trapnell et al, 2009, 2012; Kim et al, 2013). Reads per million bases and differential expression between two conditions were calculated using Cufflinks v2.2.0 (Trapnell et al, 2012). To identify differentially expressed genes, specific parameters (p-value ≤ 0.05; statistical correction: Benjamini Hochberg; FDR ≤ 0.05) in cuffdiff were used. Post-processing and visualization of differential expression were done using cummeRbund v2.0.0. A cut-off of 2 fold change and p-value less than 0.05 was set to identify the up and down regulated genes between two conditions compared. Venny (Oliveros JC 2018) was used to identify the genes common or unique to different conditions compared. AgriGO (Tian T et al, 2017) was used to find the corresponding GO terms (FDR ≤ 0.05) and the functions of the respective genes.

### Chromatin immunoprecipitation

We conducted ChIP as described in previous studies (Lamke et al, 2016a). In short, roughly 500 mg of 10 and 12 days old seedlings were cross-linked by vacuum-infiltrating 1% formaldehyde for 15 min. Formaldehyde was quenched using 2M glycine. Samples were stored at -80°C until further processing. Further, nuclei extraction was performed and chromatin was sonicated using a Diagenode Bioruptor (medium setting, 14 cycles each with 30 sec on/ 30 sec off with ice cooling), yielding fragments with a size of around 250 bp. Antibodies (anti-H3, ab1791; anti-H3K4me3, from Abcam, http://www.abcam.com) were pre-coupled to protein A-coated agarose beads (Invitrogen) for at least 2 h at 4°C. Immunoprecipitations were done in IP buffer at 4°C for overnight. After, washing and reverse crosslinking, resulted DNA was extracted using the phenol-chloroform method and precipitated with ice chilled ethanol and glycogen (Invitrogen), then re-suspended in 20 µl of water. ChIP-PCR was performed for three regions of *APX2* and two regions of *HSP18.2* gene loci. Amplification values were normalized to H3 (normalized signal modification/normalized signal H3). The given values in graphs are the means of three biological replicates, with each replicate was normalized to the respective 22°C control with no heat stress (NHS) sample before averaging.

## Acknowledgement

We would like to thank Charng YY for *hsfa2* and *hsfa1-q* mutant seeds. We thank all members of the Hirt Lab, especially Dr Naganand Rayapuram for his scientific suggestions and Olga Artyukh for her assistance at several points; CDA management team and greenhouse facility in KAUST for the technical assistance and in many other aspects.

## Author contributions

KS and HH conceptualized and designed the experiments. KS standardized the heat stress protocol and phenotyping experiments. AS performed qRT-PCR for targeted transcriptome at 1, 24 and 48 h and helped with Chip-qPCR experiment. KM performed bioinformatics analysis of raw RNA-seq data. RJ helped with phenotype experiments and RNA-extractions. KS analyzed the data. HH supervised the experimental design, execution of experiments and analysis of the data. KS wrote the manuscript, HH provided input and edited the manuscript. All authors approved the final version of the manuscript.

## Availability of data and materials

The data set supporting the results of this article is included within the article and its additional files. The raw data from the RNA-seq samples were submitted to the National Centre for Biotechnology Information (NCBI) Gene Expression Omnibus (GEO) database, under project GSE143635, accession numbers GSM4264089 - GSM4264106. The data are publicly available and accessible at https://www.ncbi.nlm.nih.gov/geo/query/acc.cgi?acc=GSE143635.

## Supplementary material

**Appendix table S1**: Summary of genes differentially up- and down-regulated in HS, HS+187, ACC and ACC+187 compared to NHS with and without SA187. List of DEGs unique to HS (HS vs. NHS) as shown in Fig. 3A-B.

**Appendix table 2:** Summary of DEGs: NHS+187 vs NHS; HS+187 vs HS; ACC vs HS

**Appendix table 3:** List of common DEGs: HS+187 vs HS and ACC vs HS.

**Appendix table 4:** List of all primers used in the study.

**Expanded view figure 1:** Histograms showing enriched GO terms for common DEGs (A) up- and (B) down-regulated genes in HS vs NHS, ACC vs NHS, HS+187 vs NHS+187, ACC+187 vs NHS+187.

**Expanded view figure 2:** Histograms showing enriched GO terms for DEGs (A) up- and (B) down-regulated genes in ACC vs HS.

**Expanded view 3:** Fresh weight of SA187-colonized or non-colonized wild type (Col-0) and *hsfa2* mutant plants under control conditions (NHS).

---

**Expanded view figure 1:**
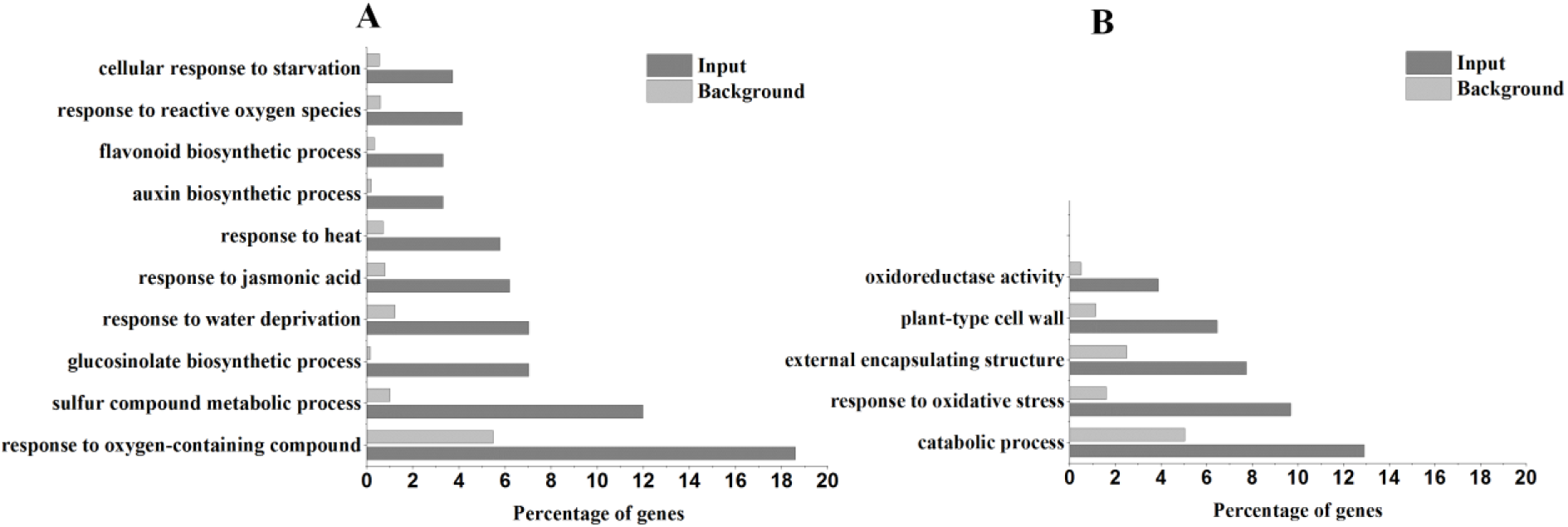
The histograms showing enriched GO terms for commonly DE up- (A) and down-regulated (B) genes in HS, ACC with +/-187 compare to NHS+/187.

**Expanded view figure 2:**
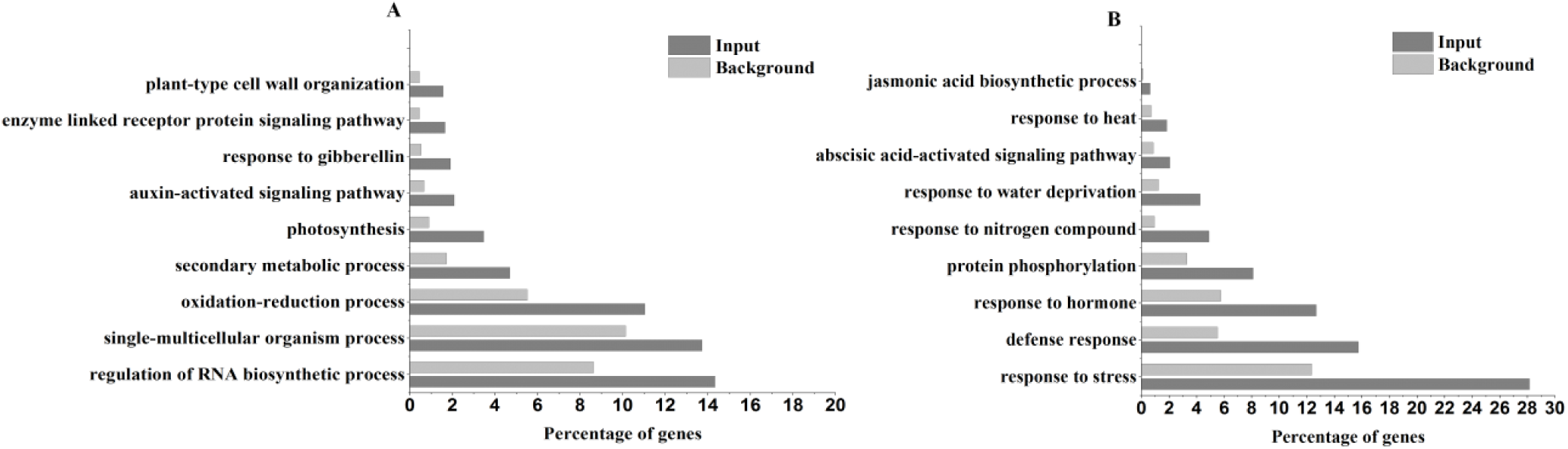
The histograms showing enriched GO terms for DE up- (A) and down-regulated (B) genes in ACC compare to HS.

**Expanded view figure 3:**
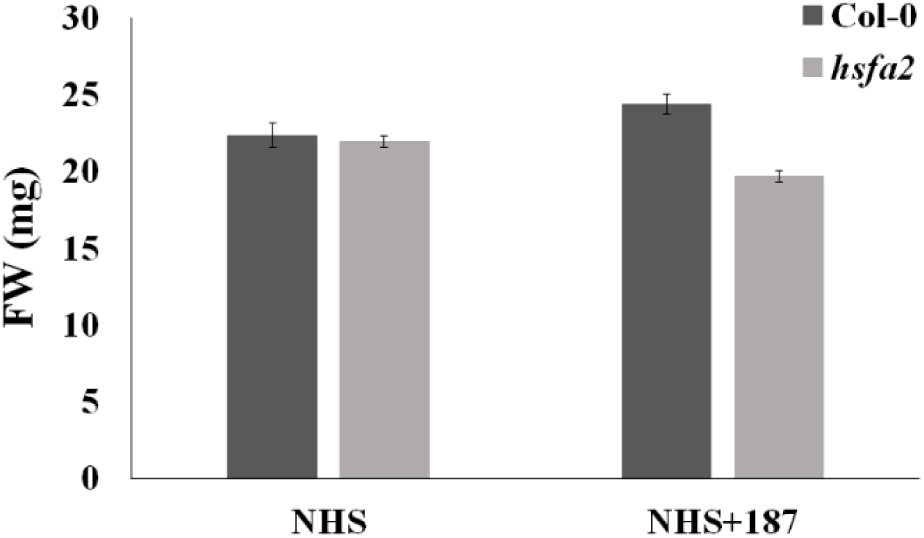
Fresh weight of SA187-colonized or non-colonized wild type (Col-0) and *hsfa2* mutant plants under control conditions (NHS).

## References

Andrés-Barrao C, Lafi FF, Alam I, de Zélicourt A, Eida AA, Bokhari A, et al. (2017). Complete Genome Sequence Analysis of Enterobacter sp. SA187 a Plant Multi-Stress Tolerance Promoting Endophytic Bacterium. Front Microbiol 8: 1–21.

Andrews S (2012). “Babraham Bioinformatics - FastQC A Quality Control tool for High Throughput Sequence Data.” (2012) (November 5, 2017).

Asseng S, Ewert F, Martre P, Rötter RP, Lobell DB, Cammarano D, et al. (2015). Rising temperatures reduce global wheat production. Nat Clim Chang 5: 143–147.

Avramova Z (2015). Transcriptional “memory” of a stress: transient chromatin and memory (epigenetic) marks at stress-response genes. Plant J 83: 149–159.

Berry S, Dean C (2015). Environmental perception and epigenetic memory: mechanistic insight through FLC. Plant J 83: 133–148.

Bolger AM, Lohse M, Usadel B, Trimmomatic (2014). A flexible trimmer for Illumina sequence data. Bioinformatics 30: 2114–20.

Brickner DG, Cajigas I, Fondufe-Mittendorf Y, Ahmed S, Lee PC, Widom J and Brickner JH (2007). H2A.Z-mediated localization of genes at the nuclear periphery confers epigenetic memory of previous transcriptional state. PLoS Biol 5: e81.

Bruce TJA, Matthes MC, Napier JA and Pickett JA (2007). Stressful “memories” of plants: evidence and possible mechanisms. Plant Sci 173: 603–608.

Brzezinka K, Altmann S, Czesnick H et al. (2016). Arabidopsis FORGETTER1 mediates stress-induced chromatin memory through nucleosome remodeling. eLIFE 5 e17061.

Cheng MC, Liao PM, Kuo WW and Lin TP (2013). The Arabidopsis ETHYLENE RESPONSE FACTOR1 regulates abiotic stress-responsive gene expression by binding to different cis-acting elements in response to different stress signals. Plant Physiol 162(3):1566–1582.

Craita B and Tom G (2013). Plant tolerance to high temperature in a changing environment: scientific fundamentals and production of heat stress-tolerant crops. Front Plant Sci 4: 273.

de Zelicourt A, Al-Yousif M, Hirt H (2013). Rhizosphere microbes as essential partners for plant stress tolerance. Mol plant 6(2): 242.

de Zélicourt A, Synek L, Saad MM, Alzubaidy H, Jalal R, Xie Y et al. (2018). Ethylene induced plant stress tolerance by Enterobacter sp. SA187 is mediated by 2-keto-4-methylthiobutyric acid production. PLOS Genet 14(3).

D’Urso A and Brickner JH (2017). Epigenetic transcriptional memory. Curr genet 63(3): 435–439.

Eida AA, Ziegler M, Lafi FF, Michell CT, Voolstra CR, Hirt H, Saad MM (2018). Desert plant bacteria reveal host influence and beneficial plant growth properties. PLOS ONE 13(12).

El-Daim I, Bejai S and Meijer J (2014). Improved heat stress tolerance of wheat seedlings by bacterial seed treatment. Plant Soil 379(1/2): 337–350.

Hilker M, Schwachtje J, Baier M, Balazadeh S, Bäurle I. Geiselhardt S, Hincha DK, Kunze R, Mueller-Roeber B, Rillig MC, Rolff J, Romeis T, Schmülling T, Steppuhn A, van Dongen J, Whitcomb SJ Wurst S, Zuther E and Kopka J (2016). Priming and memory of stress responses in organisms lacking a nervous system. Biol Rev Camb Philos Soc 91(4):1118–1133.

Kim D, et al, (2013). TopHat2: accurate alignment of transcriptomes in the presence of insertions, deletions and gene fusions. Genome Biol 14: R36.

Laine JP, Singh BN, Krishnamurthy S and Hampsey M (2009). A physiological role for gene loops in yeast. Genes Dev 23: 2604–2609.

Lamke J, Brzezinka K, Altmann S and Bäurle I (2016a). A hit-and-run heat shock factor governs sustained histone methylation and transcriptional stress memory. The EMBO J 35. 162–175.

Lamke J, Brzezinka K, Bäurle I (2016b). HSFA2 orchestrates transcriptional dynamics after heat stress in Arabidopsis thaliana. Transcription 7. 111–114.

Larkindale J and Vierling E (2008). Core genome responses involved in acclimation to high temperature. Plant Physiol 146: 748–761.

Larkindale J, Vierling E (2008). Core genome responses involved in acclimation to high temperature. Plant Physiol 146: 748–761.

Light WH, Brickner DG, Brand VR and Brickner JH (2010). Interaction of a DNA zip code with the nuclear pore complex promotes H2A.Z incorporation and INO1 transcriptional memory. Mol Cell 40: 112–125.

Lin J, Kuo C, Yang I, Tsai W, Shen Y, Lin C, Liang Y, Li Y, Kuo Y, King Y, Lai H and Jeng S (2016). MicroRNA160 Modulates Plant Development and Heat Shock Protein Gene Expression to Mediate Heat Tolerance in Arabidopsis. Front Plant Sci (9): 68.

Ling Y, Serrano N, Gao G, Atia M, Mokhtar M, Woo YH, Jeremie Bazin J, Veluchamy A, Benhamed M, Crespi M, Gehring C, Reddy ASN and Mahfouz MM (2018). Thermopriming triggers splicing memory in Arabidopsis. J exp bot 69(10): 2659–2675.

Liping X, Di Zhaocan D, Yang Wenwu Y, Liu Jiaqian L, Li Meina L, Wang Xiaojuan W, Cui Chaofan C, Wang Xiaoyun W, Wang Xiue W, Zhang Ruiqi Z, Xiao Jin X and Aizhong C (2017). Overexpression of ERF1-V from Haynaldia villosa can Enhance the Resistance of Wheat to Powdery Mildew and Increase the Tolerance to Salt and Drought Stresses. Front Plant Sci 8: 1948.

Liu H, Lämke J, Lin S, Hung M, Liu K, Charng Y and Bäurle I (2018). Distinct heat shock factors and chromatin modifications mediate the organ-autonomous transcriptional memory of heat stress. Plant J 95 (3): 401–413.

Liu HC and Charng YY (2012). Acquired thermotolerance independent of heat shock factor A1 (HsfA1). the master regulator of the heat stress response. Plant Signaling Behav 7: 547–550.

Liu HC and Charng YY (2013). Common and distinct functions of Arabidopsis class A1 and A2 heat shock factors in diverse abiotic stress responses and development. Plant Physiol 163. 276–290.

Liu HC, Liao HT and Charng YY (2011). The role of class A1 heat shock factors (HSFA1s) in response to heat and other stresses in Arabidopsis. Plant Cell Environ 34: 738–751.

Liu J, Feng L, Li J and He Z (2015). Genetic and epigenetic control of plant heat responses. Front Plant Sci 6: 267

Márquez LM, Redman RS, Rodriguez RJ and Roossinck MJ (2007). A virus in a fungus in a plant: three-way symbiosis required for thermal tolerance. Science (315): 513–515.

Mishra SK, Tripp J, Winkelhaus S, Tschiersch B, Theres K, Nover L and Scharf KD (2002). In the complex family of heat stress transcription factors, HsfA1 has a unique role as master regulator of thermotolerance in tomato. Genes Dev 16: 1555–1567.

Mittler R and Blumwald E (2010). Genetic Engineering for Modern Agriculture: Challenges and Perspectives. Annu Rev Plant Biol 61: 443–462.

Nishizawa A, Yabuta Y, Yoshida E, Maruta T, Yoshimura K and Shigeoka S (2006). Arabidopsis heat shock transcription factor A2 as a key regulator in response to several types of environmental stress. Plant J 48: 535–547.

Numan M, Bashir S, Khan Y, Mumtaz R, Khan Z, Khan AL, Khan A, and AL-Harrasi A (2018). Plant growth promoting bacteria as an alternative strategy for salt tolerance in plants: A review Microbiol Res (209): 21–32.

Oliveros JC, “Venny 2.1.0” (November 5, 2018).

Prime-A-Plant Group 2006. Priming: getting ready for battle. Mol Plant Microbe Interact 19. 1062–1071.

Sani E, Herzyk P, Perrella G, Colot V and Amtmann A (2013). Hyperosmotic priming of Arabidopsis seedlings establishes a long-term somatic memory accompanied by specific changes of the epigenome. Genome Biol 14: R59.

Schramm F, Ganguli A, Kiehlmann E, Englich G, Walch D and Koskull-Döring P (2006). The heat stress transcription factor HsfA2 serves as a regulatory amplifier of a subset of genes in the heat stress response in Arabidopsis. Plant Mol Biol 60: 759–772

Sedaghatmehr M, Mueller-Roeber B and Balazadeh S (2016). The plastid metalloprotease FtsH6 and small heat shock protein HSP21 jointly regulate thermomemory in Arabidopsis. Nat Commun. (7): 12439.

Solano R, Stepanova A, Chao Q and Ecker JR (1998). Nuclear events in ethylene signaling: a transcriptional cascade mediated by ETHYLENE-INSENSITIVE3 and ETHYLENE-RESPONSE-FACTOR1. Genes Dev. 12: 3703–3714.

Stief A, Brzezinka K, Lämke J and Bäurle I (2014). Epigenetic responses to heat stress at different time scales and the involvement of small RNAs. Plant Signaling Behav (9).

Tan-Wong SM, Wijayatilake HD andProudfoot NJ (2009). Gene loops function to maintain transcriptional memory through interaction with the nuclear pore complex. Genes Dev 23: 2610–2624.

Tian T, et al, (2017). agriGO v2.0: a GO analysis toolkit for the agricultural community, 2017 update. Nucleic Acids Res 45: W122–W129.

Trapnell C, et al, (2012). Differential gene and transcript expression analysis of RNA-seq experiments with TopHat and Cufflinks. Nat Protoc 7: 562–78.

Trapnell C, Pachter L, Salzberg SL, (2009). TopHat discovering splice junctions with RNA-Seq. Bioinformatics 25: 1105–11.

Vriet C, Hennig L and Laloi C (2015). Stress-induced chromatin changes in plants: of memories, metabolites and crop improvement. Cell Mol Life Sci 72: 1261–1273.

Wu Z, Liang J, Wang C, Zhao X, Zhong X, Cao X, Li G, He J and Yi M (2018). Overexpression of lily HsfA3s in Arabidopsis confers increased thermotolerance and salt sensitivity via alterations in proline catabolism. J Exp Bot 69(8): 2005–2021.

Yang C, Lu X, Ma B, Chen S and Zhang J (2015). Ethylene Signaling in Rice and Arabidopsis: Conserved and Diverged Aspects. Mol Plant 8 (4): 495–505.

Yeh CH, Kaplinsky NJ, Hu C and Charng YY (2012). Some like it hot some like it warm: phenotyping to explore thermotolerance diversity. Plant Sci 195: 10–23.

Yoshida T, Ohama N, Nakajima J et al, (2011). Arabidopsis HsfA1 transcription factors function as the main positive regulators in heat shockresponsive gene expression. Mol Genet Genomics. 286: 321–332.

Zhouli X, Trevor MN, Jiang H, Yanhai Y (2019). AP2/ERF Transcription Factor Regulatory Networks in Hormone and Abiotic Stress Responses in Arabidopsis. Front Plant Sci. 10: 228.

